# Development of a Stable Human iPSC Line from Peripheral Blood: A Control Resource for Disease Modeling

**DOI:** 10.1101/2025.08.21.671443

**Authors:** Surajit Malakar, Vasanth Thamodaran, Tamali Halder, Deepika Joshi, Parimal Das

## Abstract

**Introduction:** Traditional disease modeling approaches have primarily utilized animal models and immortalized cell lines to investigate disease mechanisms and develop therapeutic strategies. However, previous research indicates that only about 5% of therapeutic interventions tested in animal models eventually receive regulatory approval for human use, highlighting limitations. The discovery of human-induced pluripotent stem cells (iPSCs) by Yamanaka’s team in 2007 has revolutionized the field with remarkable possibilities for modeling human diseases, drug testing, and regenerative medicine. Differentiated cells derived from iPSCs in two-dimensional (2D) monolayers offer a relatively straightforward system to study disease pathogenesis and underlying molecular mechanisms.

**Objective:** The present study aimed to generate and characterize an induced pluripotent stem cell (iPSC) line from peripheral blood mono-nuclear cells (PBMCs) of a healthy individual intending to serve as an age and gender-matched control for future disease modeling and regenerative medicine research.

**Materials and Methods:** PBMCs were isolated from a healthy 31-year-old male volunteer. Somatic reprogramming was performed using episomal vectors expressing *OCT3/4, SOX2, KLF4*, and *L-MYC*. The resulting colonies were cultured and characterized for pluripotency markers by immunocytochemistry demonstrating the ability to differentiate into three germ layers. Karyotype analysis was performed to check the chromosomal abnormalities.

**Inferences:** The resulting iPSC line exhibited pluripotency markers with the differentiation ability into three germ layers. Karyotyping analysis confirmed a normal chromosomal profile in both the donor and the reprogrammed iPSC line. This iPSC line could be a valuable resource as a healthy control for disease modeling and will contribute to advancing stem cell research with potential for regenerative medicine applications.

## 1. Introduction

The major component in unravelling the aetiology and pathophysiology of any disease mechanism, as well as in drug discovery, is the availability of physiologically relevant experimental models, either *in vivo, in vitro*, or both. While *in vivo* models have significantly contributed to basic research and therapeutic approaches, their translational success in human drug trials is often limited due to species-specific differences in biological responses [1].

The discovery of human-induced pluripotent stem cells (iPSCs) has created exciting opportunities for *in vitro* modeling of human biology and for developing cell-based therapies [2, 3]. To explore intricate disease mechanisms and misregulated protein functions, it is essential to establish a stable control line from a healthy individual [4, 5]. Traditional model animals, such as mice and rats, frequently fail to accurately represent human disease states because of anatomical differences and a lack of conservation in specific genes of interest [6].

Consequently, iPSCs derived from human donors have become increasingly vital in research. These cells not only provide a more relevant platform for studying human diseases but also enable the development of personalized therapies tailored to individual genetic backgrounds [7]. In addition, iPSCs support high-throughput screening of drug candidates, thereby accelerating the identification of effective treatments [8]. With continued refinement of these technologies, iPSCs hold great promise for advancing regenerative medicine and disease modeling, ultimately improving patient outcomes [9].

In this context the present study has been designed to generate a control iPSC line from a healthy individual using non-integrative episomal vector expressing Yamanaka factors. This reprogrammed iPSC line exhibits embryonic stem cell like colonies and robust expression of pluripotent markers.

This newly produced iPSC line is an excellent resource for studying biological processes, disease etiology, and therapeutic applications [10]. Its production contributes to the growing pool of patient- and control-derived iPSCs, easing future attempts in personalized medicine and functional genomics [7].

## 2. Materials and Methods

### 2.1. Volunteer Selection

A healthy 31-year-old male volunteer was recruited for this study after a complete medical examination at the Institute of Medical Sciences (IMS), Banaras Hindu University (BHU). The stem cell culture facility at TIGS, Bangalore, was utilized for the generation of iPSCs. Prior to reprogramming, the subject’s blood was screened for hepatitis B surface antigen (HBsAg) using a rapid test and for HIV-1 and HIV-2 antibodies at NABL M(EL)T Labs, Bangalore, India.

### 2.2. Isolation of PBMCs

5 ml of peripheral blood was collected from the subject for PBMC isolation using Ficoll-Paque Premium, a Ficoll-based density gradient medium. An equal volume of peripheral blood, diluted with an equal volume of Cell Therapy Systems Dulbecco’s Phosphate Buffered Saline (CTS-DPBS), was carefully layered on top of the density gradient medium and centrifuged at 400 × g for 36 minutes at 18–20°C. The PBMCs located at the interphase were carefully collected into a new 15 ml tube and treated with RBC lysis buffer. Following lysis, the PBMCs were washed twice with CTS DPBS, frozen in FBS containing 20% DMSO at a concentration of 5 × 10^6^ cells/ml, and stored in liquid nitrogen until further use [11].

### 2.3. Reprogramming PBMCs to iPSCs

PBMCs at a density of 5 × 10^6^ cells/ml were cultured in SFEM II medium (STEMCELL technologies, 09605) supplemented with growth factors SCF (50 ng/ml), IGF (40 µg/ml), EPO (3 U/ml), and IL-3 (10 µg/ml). On day 8, PBMCs were nucleofected with three episomal plasmids (pCXLE-hOCT3/4-shp53-F, pCXLE-hSK, and pCXLE-hUL) encoding pluripotency factors OCT4 with shRNA against p53, SOX2, KLF4, L-MYC, and LIN28 (Addgene, #27077, #27078, #27080), in a 1:1:2 ratios. Nucleofection was performed with 0.5 million cells using the Invitrogen Neon Transfection System under the following conditions: 1300 V, 30 ms pulse width, and 1 pulse. Nucleofected cells were cultured for 22 days at 37°C with 5% CO2 in 35 mm plates coated with 1% Geltrex (Thermo, A1413301) and supplemented with 2 ml of SFEM II medium containing growth factors (SCF, IGF, EPO, and IL-3). From days 3 to 5, the medium was gradually replaced with StemFlex medium (Gibco, A33492-01) until the complete switch was achieved [11].

### 2.4. iPSC Maintenance

iPSCs were maintained on 1% Geltrex-coated plates in StemFlex complete medium, with medium changes every other day. Colonies free of differentiation were manually picked or dissociated with 1× EDTA and sub-cultured onto new Geltrex-coated plates.

### 2.5. Immunofluorescence

iPSCs cultured in 24-well plates were fixed with 4% paraformaldehyde for 20 minutes. Cells were then permeabilized with 0.2% Triton X-100 (sigma) for 10 minutes at room temperature and washed once with PBS. Fixed cells were incubated overnight at 4°C with primary antibodies, followed by incubation with secondary antibodies for 2 hours at room temperature. DAPI was added for nuclear staining and incubated with gentle shaking for 15 minutes. Cells were then washed with PBS before observation **[Table 2]**.

### 2.6. Karyotyping

GTG banding analysis was performed at Neuberg Anand Diagnostic Laboratory, Bangalore, to assess chromosomal integrity and exclude aneuploidy.

### 2.7. Tri-Lineage Differentiation

#### 2.7.1. Endoderm differentiation

iPSCs were seeded at a density of 2.1 × 10^6^ cells/cm^2^ on 1% Geltrex-coated plates in StemFlex medium for 48 hours. Cells were then cultured for 48 hours in endoderm differentiation medium consisting of RPMI 1640 (Thermo, 11875093), 0.2% FBS, 25 ng/ml Activin A, 2 µM CHIR 99021, 1× NEAA, and 5 mM crotonate at pH 7.4. Differentiation was confirmed by SOX17 immunostaining [12].

#### 2.7.2. Mesoderm differentiation

iPSCs dissociated with TrypLE (Thermo, 12604021) were seeded at 2.1 × 10^6^ cells/cm^2^ on 1% Geltrex-coated plates and maintained in StemFlex medium for 48 hours. Cells were then cultured in mesoderm differentiation medium consisting of Advanced RPMI (Thermo, 12633012), GlutaMAX (Thermo, 35050061), and 8 µM CHIR. Differentiation was confirmed by Brachyury immunostaining.

#### 2.7.3. Ectoderm differentiation

iPSCs (4 × 10^6^ cells/well) were seeded on 1% Geltrex-precoated 24-well plates and cultured for 6 days in ectodermal differentiation medium I, composed of DF12: Neurobasal medium (Thermo, 21103049), GlutaMAX, 3 µM CHIR, 0.1% B27 (with insulin), 0.1 mM ascorbic acid, 2 µM DMH1, 2 µM SB431542, and 0.5% N2. After 6 days, cells were dissociated and reseeded in ectodermal differentiation medium II, containing DF12: Neurobasal medium, GlutaMAX, 1 µM CHIR, 0.1% B27 (with insulin), 0.1 mM ascorbic acid, 2 µM DMH1, 2 µM SB431542, 0.5% N2, 0.5 µM puromycin, and 0.1 µM retinoic acid, for an additional 6 days. Differentiation was confirmed by PAX6 immunostaining [13].

#### 2.8. Testing for Mycoplasma contamination

600 µL of iPSC spent media was used for DNA isolation using equal volumes of TE-saturated phenol and 3M sodium acetate, followed by PCR amplification using primers of the conserved region in the mycoplasma 16S RNA gene.

## 3. Result

## 3.1. PBMC Isolation and iPSC Derivation

PBMC was isolated successfully from subject’s peripheral blood using ficoll based density gradient centrifugation [11], and cultured for 8 days. Reprogramming into iPSCs was performed using an episomal vector system encoding the Yamanaka factors (OCT4, SOX2, KLF4, and L-MYC) **[Fig 1 A, B]**.

**Fig 1:**
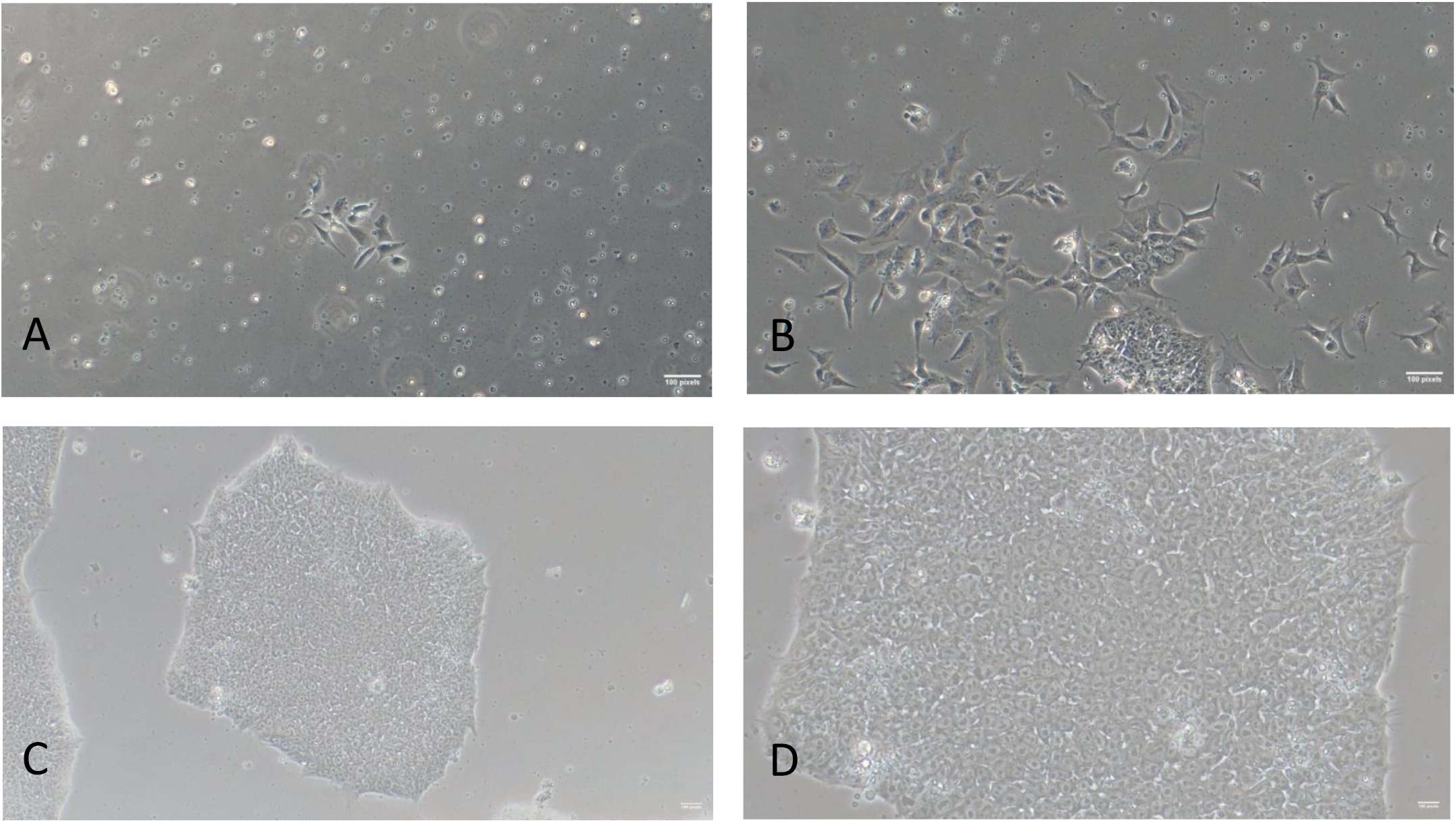
iPSC reprogramming observed by Phase-contrast microscopy: A. Day 9 10X B. Day15 10X C. Day 20 10X D. Day 20 20X.

### 3.2. Morphology

By day 19 of post-nucleofection, compact, dome-shaped colonies with high nuclear-to-cytoplasmic ratios and prominent nucleoli were observed, consistent with human embryonic stem cell morphology **[Fig 1 C, D]**.

### 3.3. Pluripotency

Pluripotency was confirmed by Immunocytochemistry through expression of NANOG, SOX2, OCT4, TRA-1-60, TRA-1-81, and SSEA4 **[Fig 2] [Table 1]**.

**Table 1:** List of tests used for Characterization of iPSC.

| Classification | Test | Result | Data |
| --- | --- | --- | --- |
| Morphology | Phase contrast microscopy | Healthy and compact cell colonies with defined borders resembles embryonic cells | Fig 1 A,B,C,D |
| Phenotype | Immunostaining | Expression of pluripotent markers, OCT4, SOX2, NANOG, TRA1-61, TRA-181, SSEA4 | Fig 2 |
| Genotype | Karyotyping (G banding) | No chromosomal abnormalities, 46XY | Fig 3 |
| Differentiation | Trilineage | Expression of | Fig 4 |
| potential | differentiation,<br>immunostaining | Endoderm- SOX17,<br>Mesoderm-BRACHYURY and<br>Ectoderm- PAX6 |  |
| Microbiology and<br>Virology | Mycoplasma test | Sample tested<br>negative | Fig 5 |
| Donor screening | HIV-1+2, Hepatitis<br>B | Negative |  |

**Table 2:** List of antibodies used in the study.

| **Marker** | **Antibody** | **Dilution** | **Company**<br>**Catalogue** |
| --- | --- | --- | --- |
| Pluripotent marker | OCT4 Rabbit mAb | 1:1000 | Cell signaling<br>technology, 9656s |
| Pluripotent marker | SSEA4mousemAb | 1:1000 | Cell signaling<br>technology, 9656s |
| Pluripotent marker | SOX2 Rabbit mAb | 1:1000 | Cell signaling<br>technology, 9656s |
| Pluripotent marker | NANOG Rabbit mAb | 1:1000 | Cell signaling<br>technology, 9656s |
| Pluripotent marker | TRA-1-60 mousemAb | 1:1000 | Cell signaling<br>technology, 9656s |
| Pluripotent marker | TRA-1-81 mousemAb | 1:1000 | Cell signaling<br>technology, 9656s |
| Endoderm<br>differentiation | SOX17 Rabbit mAb | 1:1000 | Cell signaling<br>technology, #81778 |
| Mesoderm<br>differentiation | T- Brachyury Rabbit<br>mAb | 1:1000 | Cell signaling<br>technology, #81694 |
| Ectoderm<br>differentiation | PAX6 Rabbit mAb | 1:1000 | Cell signaling<br>technology, # 60433 |
| Secondary antibody | Alexa 555 goat anti-<br>rabbit | 1:1000 | Invitrogen, A-21428 |
| Secondary antibody | Alexa 488 goat anti-<br>mouse | 1:1000 | Invitrogen, A-11001 |

**Fig 2:**
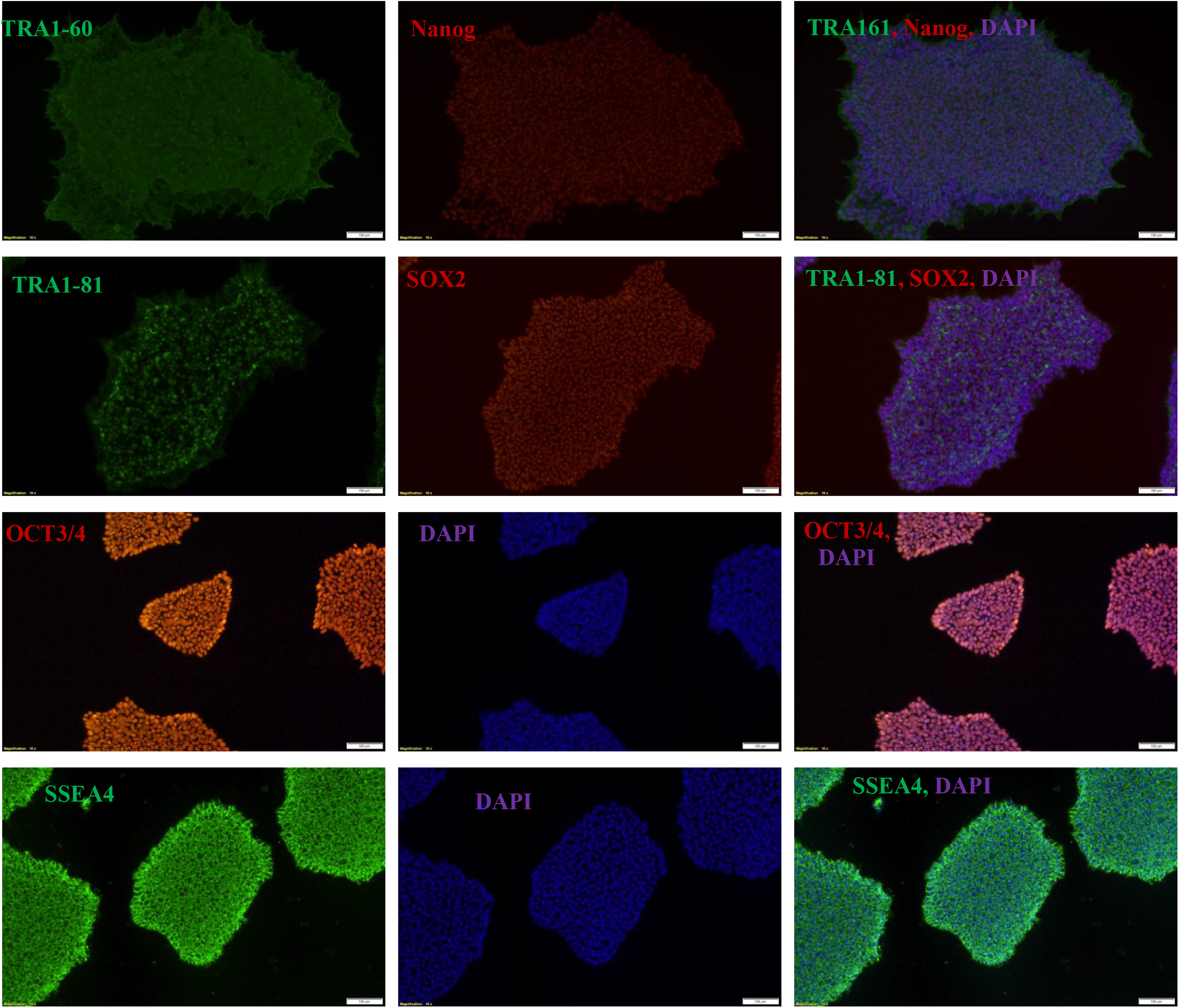
Expression of Pluripotency markers observed by fluorescence microscopy.

### 3.4. Karyotype

Karyotyping analysis at passage 20 revealed a normal male chromosomal complement (46, XY) without detectable anomalies **[Fig 3]**.

**Fig 3:**
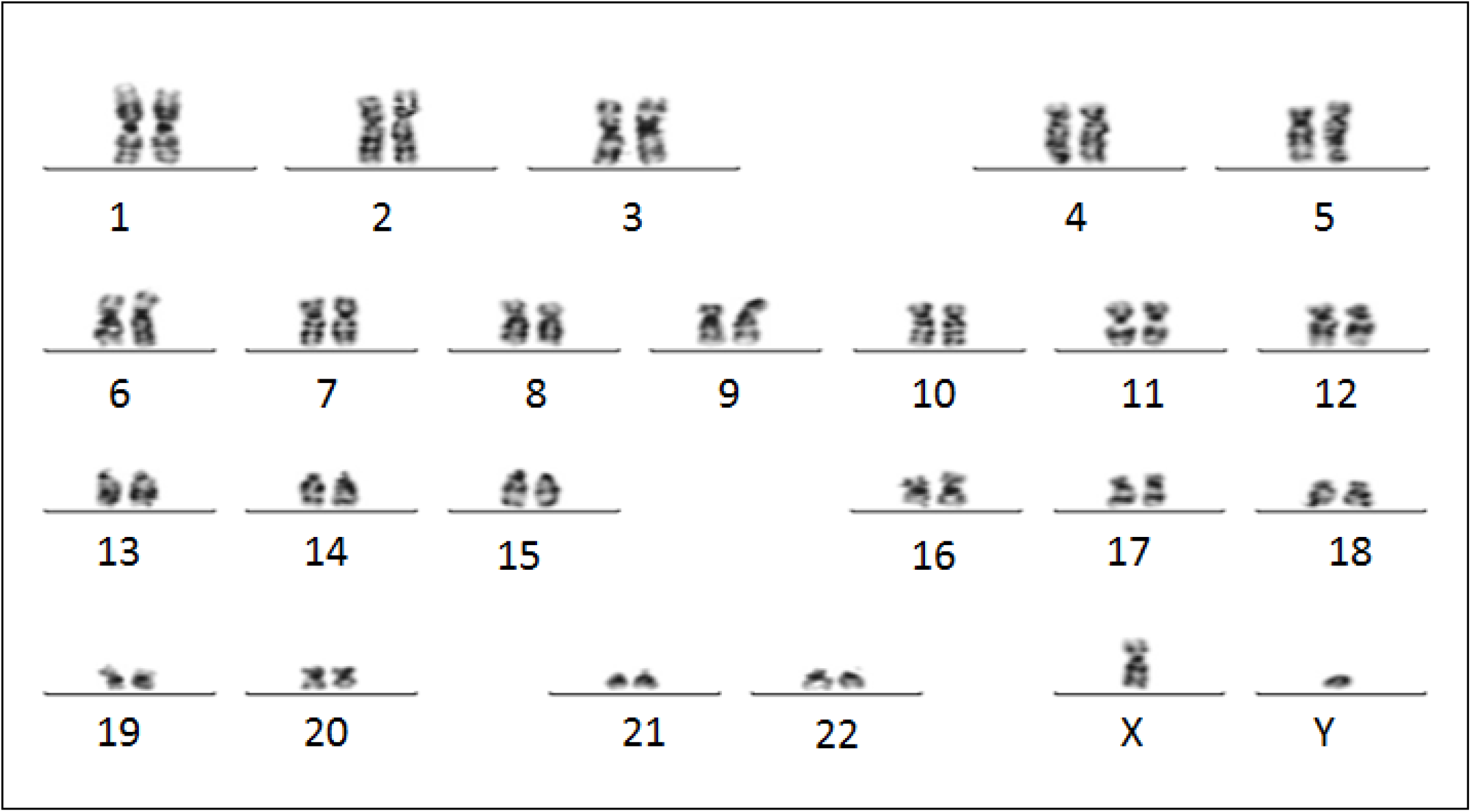
Karyotyping of iPSC.

### 3.5. Differentiation

The expression of SOX17 (endoderm), BRACHYURY (mesoderm), and PAX6 (ectoderm), were validated by immunofluorescence, demonstrating the ability of iPSCs to differentiate into all three germ layers **[Fig 4] [Table 1]**.

**Fig 4:**
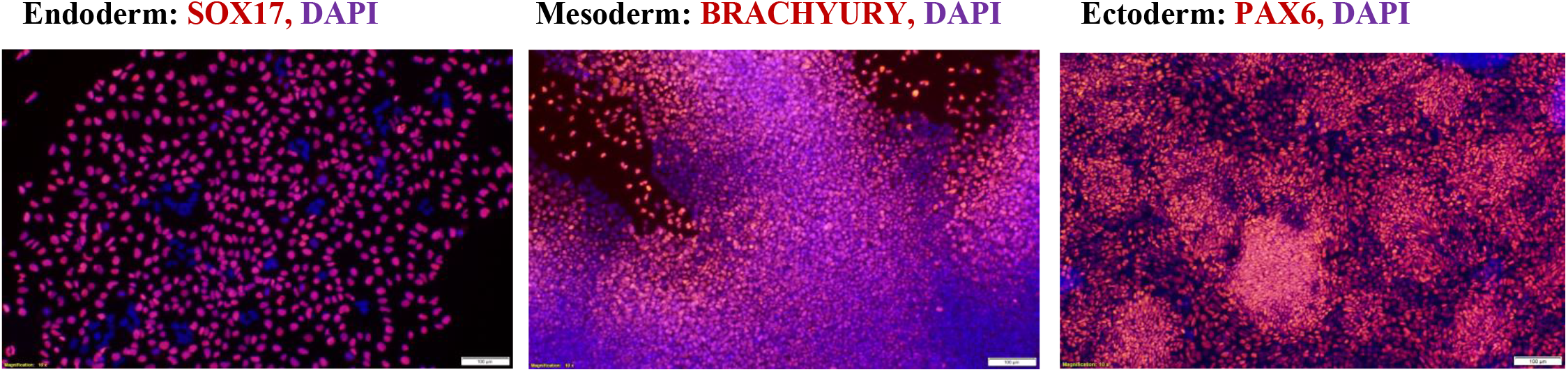
*In vitro* tri-lineage differentiation observed by fluorescence microscope.

### 3.6. Contamination-Free Status

No mycoplasma band of 800bp was observed in the established iPSC line **[Fig 5]**.

**Fig 5:**
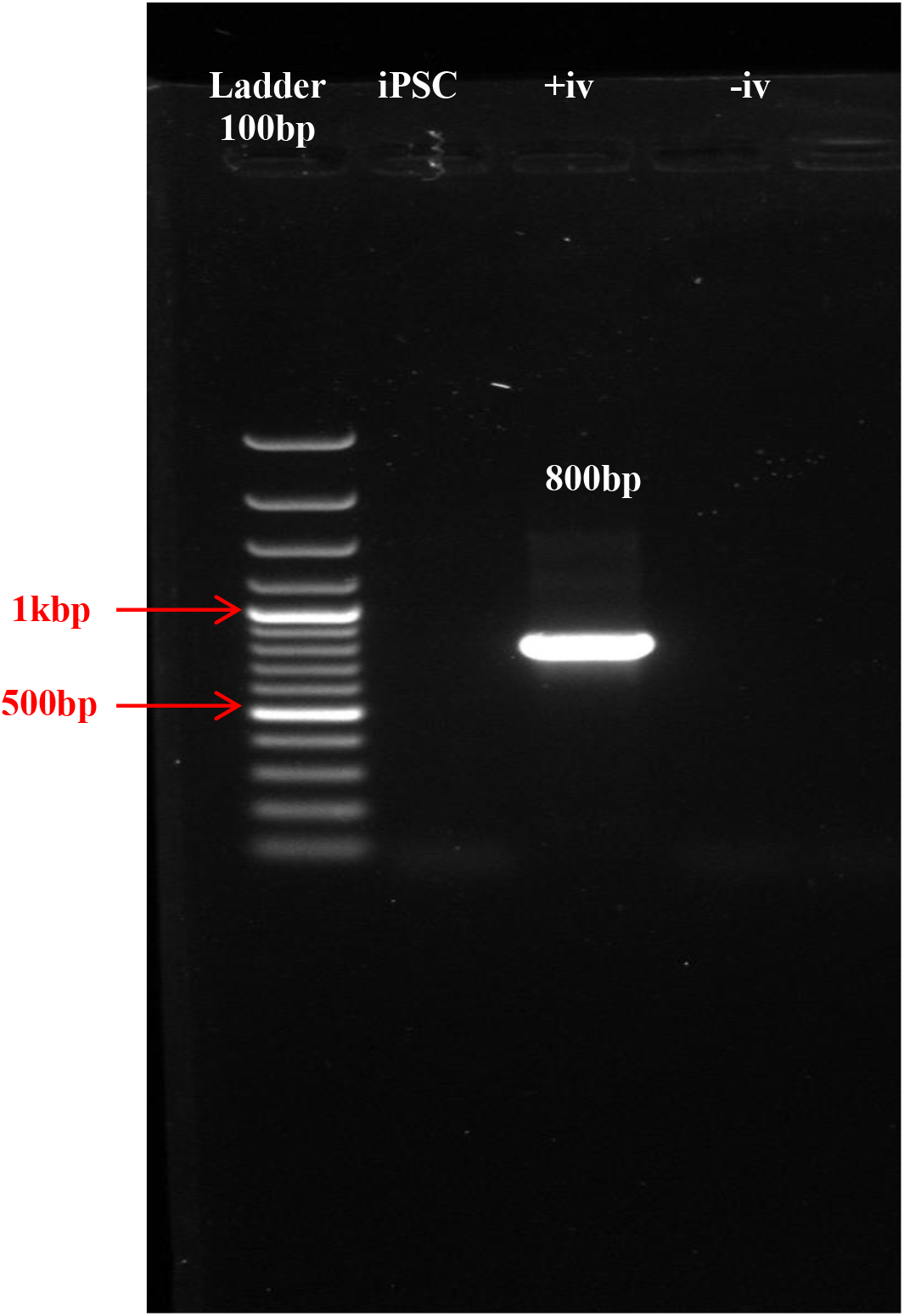
Representative gel image showing mycoplasma contamination status.

## 4. Discussion

The use of a non-integrating episomal vector approach minimized genomic integration risks, thereby enhancing the safety profile of the generated lines for potential downstream applications. Previous studies have shown that non-integrating methods produce iPSCs with fewer genomic aberrations and improved suitability for clinical translation compared to integrating viral systems [14, 15]. Moreover, the high viability and normal karyotype observed across passages underscore the stability of the reprogrammed cells, in line with earlier findings [16].

In this study, a hiPSC line was developed from PBMCs of a healthy individual. The PBMCs were reprogrammed using episomal vectors expressing OCT3/4, SOX2, KLF4, and L-Myc. The reprogrammed cells exhibited embryonic stem cell-like colonies with distinct boundaries and a high nuclear-to-cytoplasm ratio [17, 18]. Pluripotency markers including OCT4, SOX2, NANOG, TRA-1-60, TRA-1-81, and SSEA4 were confirmed by immunofluorescence **[Fig 2] [Table 1]**. The line was free of mycoplasma contamination and chromosomal abnormalities. The established iPSC line demonstrated the potential to differentiate into all three germ layers— endoderm, mesoderm, and ectoderm—via spontaneous differentiation with small molecules, confirmed by the expression of SOX17, BRACHYURY, and PAX6, respectively [19] **[Fig 4] [Table 1]**.

Although the efficiency of reprogramming remains relatively low compared to direct lineage conversion, the yield obtained in this study was within the expected range for similar protocols [20]. Several factors, including donor cell type, passage number, and culture conditions, are known to influence reprogramming efficiency [21]. Future optimization through the use of small-molecule enhancers [22] or alternative factor combinations [23] could further improve reprogramming outcomes and reduce variability between cell lines.

Overall, our findings reinforce the feasibility of generating stable and pluripotent iPSCs from somatic cells. These lines not only provide a renewable source of patient-specific stem cells but also establish a robust foundation for downstream differentiation studies, disease modeling, and potential applications in regenerative medicine [24].

## 5. Conclusion

In this study, human somatic cells were successfully reprogrammed into induced pluripotent stem cells (iPSCs) using an non-integrating episomal vector approach. The resulting colonies displayed typical stem cell morphology, expressed key pluripotency markers, and maintained normal karyotype stability. These findings confirm the efficiency and reliability of the reprogramming process and provide a robust platform for generating patient-specific iPSC lines. The established reprogramming protocol can serve as a foundation for downstream applications in disease modeling, drug screening, and regenerative medicine.

## 7. Acknowledgement

The laboratory facility, TIGS; stem cell lab members, TIGS, Bangalore is acknowledged for the present study.

## References

1. Doss, M. X., & Sachinidis, A. (2019). Current challenges of iPSC-based disease modeling and therapeutic implications. Cells, 8(5), 403. 10.3390/cells8050403

2. Wang, A. Y. L., Aviña, A. E., Liu, Y. Y., & Kao, H. K. (2025). Pluripotent stem cells: Recent advances and emerging trends. Biomedicines, 13(4), 765. 10.3390/biomedicines13040765

3. Wang, J., Huang, S., Li, K., Li, J., & Huang, X. (2025). Global research dynamics in the induced pluripotent stem cell and diabetes: A bibliometric analysis of the past twenty years. Regenerative Therapy, 30, 341–350. 10.1016/j.reth.2025.06.015

4. Brooks, I. R., Garrone, C. M., Kerins, C., Kiar, C. S., Syntaka, S., Xu, J. Z., Spagnoli, F. M., & Watt, F. M. (2022). Functional genomics and the future of iPSCs in disease modeling. Stem Cell Reports, 17(5), 1033–1047. 10.1016/j.stemcr.2022.03.019

5. Méjécase, C., Harding, P., Sarkar, H., Eintracht, J., Lima Cunha, D., Toualbi, L., & Moosajee, M. (2020). Generation of two human control iPS cell lines (UCLi016-A and UCLi017-A) from healthy donors with no known ocular conditions. Stem Cell Research, 49, 102113. 10.1016/j.scr.2020.102113

6. Reilly, L., Munawar, S., Zhang, J., Crone, W. C., & Eckhardt, L. L. (2022). Challenges and innovation: Disease modeling using human-induced pluripotent stem cell-derived cardiomyocytes. Frontiers in Cardiovascular Medicine, 9, 966094. 10.3389/fcvm.2022.966094

7. Paik, D. T., Chandy, M., & Wu, J. C. (2020). Patient and disease-specific induced pluripotent stem cells for discovery of personalized cardiovascular drugs and therapeutics. Pharmacological Reviews, 72(1), 320–342. 10.1124/pr.116.013003

8. Cerneckis, J., Cai, H., & Shi, Y. (2024). Induced pluripotent stem cells (iPSCs): Molecular mechanisms of induction and applications. Signal Transduction and Targeted Therapy, 9, 112. 10.1038/s41392-024-01809-0

9. Wiegand, C., & Banerjee, I. (2019). Recent advances in the applications of iPSC technology. Current Opinion in Biotechnology, 60, 250–258. 10.1016/j.copbio.2019.05.011

10. Saito, M. K., Osawa, M., Tsuchida, N., Shiraishi, K., Niwa, A., Woltjen, K., Asaka, I., Ogata, K., Ito, S., Kobayashi, S., & Yamanaka, S. (2023). A disease-specific iPS cell resource for studying rare and intractable diseases. Inflammation and Regeneration, 43(1), 43. 10.1186/s41232-023-00294-2

11. Singh, G., Manian, K. V., Premkumar, C., Srivastava, A., Daniel, D., & Velayudhan, S. R. (2022). Derivation of clinical-grade induced pluripotent stem cell lines from erythroid progenitor cells in xenofree conditions. Methods in Molecular Biology, 2454, 775–789. 10.1007/7651_2021_349

12. Fang, Y., & Li, X. (2021). A simple, efficient, and reliable endoderm differentiation protocol for human embryonic stem cells using crotonate. STAR Protocols, 2(3), 100659. 10.1016/j.xpro.2021.100659

13. Du, Z. W., Chen, H., Liu, H., Lu, J., Qian, K., Huang, C. L., Zhong, X., Fan, F., & Zhang, S. C. (2015). Generation and expansion of highly pure motor neuron progenitors from human pluripotent stem cells. Nature Communications, 6, 6626. 10.1038/ncomms7626

14. Takahashi, K., & Yamanaka, S. (2006). Induction of pluripotent stem cells from mouse embryonic and adult fibroblast cultures by defined factors. Cell, 126(4), 663–676. 10.1016/j.cell.2006.07.024

15. Yu, J., Vodyanik, M. A., Smuga-Otto, K., Antosiewicz-Bourget, J., Frane, J. L., Tian, S., Stewart, R., Slukvin, I. I., & Thomson, J. A. (2007). Induced pluripotent stem cell lines derived from human somatic cells. Science, 318(5858), 1917–1920. 10.1126/science.1151526

16. Okita, K., Matsumura, Y., Sato, Y., Okada, A., Morizane, A., Okamoto, S., Hong, H., Nakagawa, M., Tanabe, K., Tezuka, K., Shibata, T., Kunisada, T., Takahashi, M., Takahashi, J., Saji, H., & Yamanaka, S. (2011). A more efficient method to generate integration-free human iPS cells. Nature Methods, 8(5), 409–412. 10.1038/nmeth.1591

17. Fusaki, N., Ban, H., Nishiyama, A., Saeki, K., & Hasegawa, M. (2009). Efficient induction of transgene-free human pluripotent stem cells using a vector based on Sendai virus, an RNA virus that does not integrate into the host genome. Proceedings of the Japan Academy, Series B, Physical and Biological Sciences, 85(8), 348–362. 10.2183/pjab.85.348

18. Schlaeger, T. M., Daheron, L., Brickler, T. R., Entwisle, S., Chan, K., Cianci, A., DeVine, A., Ettenger, A., Fitzgerald, K., Godfrey, M., Gupta, D., McPherson, J., Malwadkar, P., Gupta, M., Bell, B., Dibb, N. J., McNicol, A. M., Patel, H., Patel, D. J., Daley, G. Q. (2015). A comparison of non-integrating reprogramming methods. Nature Biotechnology, 33(1), 58–63. 10.1038/nbt.3070

19. Zhou, W., & Freed, C. R. (2009). Adenoviral gene delivery can reprogram human fibroblasts to induced pluripotent stem cells. Stem Cells, 27(11), 2667–2674. 10.1002/stem.201

20. Hussein, S. M., Batada, N. N., Vuoristo, S., Ching, R. W., Autio, R., Närvä, E., Nagy, A. (2011). Copy number variation and selection during reprogramming to pluripotency. Nature, 471(7336), 58–62. 10.1038/nature09871

21. González, F., Boué, S., & Izpisua Belmonte, J. C. (2011). Methods for making induced pluripotent stem cells: Reprogramming à la carte. Nature Reviews Genetics, 12(4), 231–242. 10.1038/nrg2937

22. Li, W., Wei, W., Zhu, S., Zhu, J., Shi, Y., Lin, T., Hao, E., Hayek, A., Deng, H., & Ding, S. (2009). Generation of rat and human induced pluripotent stem cells by combining genetic reprogramming and chemical inhibitors. Cell Stem Cell, 4(1), 16–19. 10.1016/j.stem.2008.11.014

23. Warren, L., Manos, P. D., Ahfeldt, T., Loh, Y. H., Li, H., Lau, F., Ebina, W., Mandal, P. K., Smith, Z. D., Meissner, A., Daley, G. Q., Brack, A. S., Collins, J. J., Cowan, C., Schlaeger, T. M., & Rossi, D. J. (2010). Highly efficient reprogramming to pluripotency and directed differentiation of human cells with synthetic modified mRNA. Cell Stem Cell, 7(5), 618–630. 10.1016/j.stem.2010.08.012

24. Rowe, R. G., & Daley, G. Q. (2019). Induced pluripotent stem cells in disease modelling and drug discovery. Nature Reviews Genetics, 20(7), 377–388. 10.1038/s41576-019-0100-z

